# Enhanced assembly of bacteriophage T7 produced in cell-free reactions under simulated microgravity

**DOI:** 10.1101/2022.12.16.520761

**Authors:** François-Xavier Lehr, Bruno Pavletić, Timo Glatter, Thomas Heimerl, Ralf Moeller, Henrike Niederholtmeyer

**Affiliations:** Max Planck Institute for Terrestrial Microbiology, Marburg, Germany; Center for Synthetic Microbiology (SYNMIKRO), Philipps-Universität Marburg, Marburg, Germany; German Aerospace Center, Institute of Aerospace Medicine, Aerospace Microbiology, Köln/Cologne, Germany; Campus Straubing for Biotechnology and Sustainability, Technical University of Munich, Germany

**Keywords:** cell-free synthesis, clinorotation, virus assembly, microgravity, synthetic biology, spaceflight

## Abstract

On-demand biomanufacturing has the potential to improve healthcare and self-sufficiency during space missions. Cell-free transcription and translation reactions combined with DNA blueprints can produce promising therapeutics like bacteriophages and virus-like particles. However, how space conditions affect the synthesis and self-assembly of such complex multi-protein structures is unknown. Here, we characterize the cell-free production of infectious bacteriophage T7 virions under simulated microgravity. Rotation in a 2D-clinostat increased the number of infectious particles compared to static controls. Quantitative analyses by mass spectrometry, immuno-dot-blot and real-time PCR showed no significant differences in protein and DNA contents, suggesting enhanced self-assembly of T7 phages in simulated microgravity. While the effects of genuine space conditions on the cell-free synthesis and assembly of bacteriophages remain to be investigated, our findings support the vision of a cell-free synthesis-enabled “astropharmacy”.

## Introduction

Long-term space missions will put astronaut health at immense risk^1^. Healthcare options will be limited by payload constraints and drug stability. Additionally, close quarters, radiation, and microgravity strain human health, for example by compromising the immune system^2^. On-demand production of therapeutics presents a solution to adapt medical care to the special challenges of space travel^3^. This vision of an “astropharmacy” could be realized by onboarding light weight cell-free transcription-translation (TXTL) reagents capable of rapidly synthesizing RNA and proteins from DNA blueprints, just as required. Relying only on isolated biochemical components that can be freeze-dried for storage, TXTL systems combine simplicity and flexibility, while reducing the need for downstream processing^4–6^ and biocontainment in planetary protection efforts^7^. TXTL technology enables the synthesis of large, self-assembling macromolecular complexes such as bacteriophages^8,9^ and virus-like particles^10^. Bacteriophages, as viruses that specifically target bacteria, are promising tools in the fight against multi-resistant bacteria^8,11,12^. Additionally, engineered virus-like particles could be used for gene therapy, drug delivery, and other personalized therapeutic applications^13^.

Many biological processes as well as biomolecular self-assembly depend on gravity. For example, in microgravity, proteins form larger crystals with fewer defects^14^, and amyloid fibrils nucleate and grow differently in simulated microgravity^15,16^. Comparing virus assembly in orbiter flight studies against ground controls, polyomavirus assembled into larger and more homogeneous capsomeres but did not form capsid-like structures^17^. It is hypothesized that the altered and often improved self-assembly in the absence of gravity is due to abolished sedimentation and changes in molecular transport from a convection-dominated into a diffusion-dominated regime^18^. While microgravity is known to alter gene expression in microorganisms^19^ and to make DNA polymerase more error-prone *in vitro*^20^, its effects on cell-free transcription and translation remain unexplored.

Here we focus on the cell-free production of the model bacteriophage T7 under simulated microgravity (s-μg). 2D-clinorotation approximates microgravity by rotating a sample to prevent sedimentation and has been used to simulate microgravity in experiments with different cell types and organisms^21–24^. Bacteriophage T7 infects *Escherichia coli* bacteria and has a 40 kb genome, encoding 56 proteins. Each infectious phage particle is assembled from a total number of roughly 500 protein subunits and a DNA molecule that is tightly packed into the capsid^25^. In a synthesis reaction containing purified T7 DNA and TXTL reagents, the number of infectious phage particles produced depends on three major processes: transcription, translation, and self-assembly (Fig. 1). By quantifying the effects of simulated microgravity on cell-free synthesis and self-assembly of a complex multi-protein structure, our work takes a first step in assessing the possibility of TXTL-assisted on-demand manufacturing of biologics in space.

**Figure 1.**
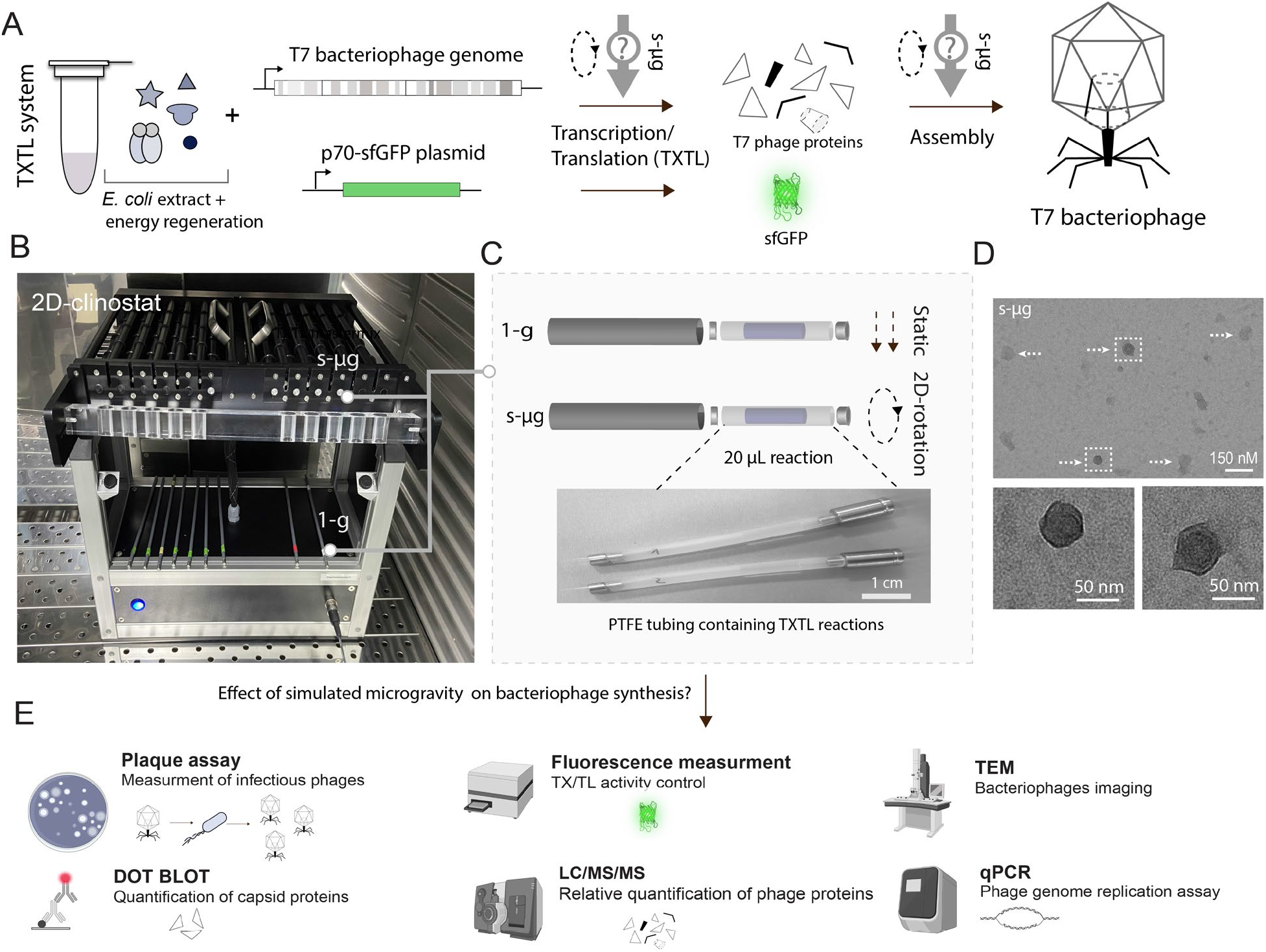
Experimental pipeline for testing the effect of simulated microgravity on the synthesis and assembly of bacteriophages in transcription-translation (TXTL) systems. A) Phage DNA is mixed with the TXTL components to start the reaction to produce bacteriophage T7. B) A 2D-clinostat is used to simulate microgravity (s-μg). C) TXTL reactions are performed in Polytetrafluoroethylene (PTFE) tubes fitted to the clinostat. Control reactions are performed in identical reaction tubes held static (1-g). D) Transmission electron microscopy (TEM) of T7 phages from the s-μg condition. Arrows indicate full size bacteriophage particles; boxes indicate phages enlarged below. E) Quantitative methods are used to assay protein and DNA content in TXTL samples (Dot-blot, mass spectrometry, fluorescence measurements, real-time quantitative PCR) and the assembly of infectious bacteriophages (plaque assays, TEM).

## Results and discussions

To accommodate TXTL reactions in the 2D-clinostat, we customized vessels to support the small volumes (10-100 μL) typically used in cell-free protein synthesis. Briefly, polytetrafluoroethylene (PTFE) tubing was enclosed into polyvinyl chloride (PVC) tubes fitting the clinostat rotational tray (Fig. 1B-C). We first verified synthesis and assembly of T7 bacteriophages were possible and reproducible in the PTFE vessels. Compared to standard microreaction tubes, we observed a slight decrease of plaque forming units (PFU) per mL, likely due to the difference in oxygen availability (Supplementary Fig. S1). Nevertheless, the low variability observed in our setup enabled us to investigate the phage synthesis kinetics in clinorotation conditions. A TXTL mastermix containing 2 nM of T7 DNA was split into samples for timepoints for s-μg and stationary control (1-g) conditions. For each timepoint, plaque assays were performed in duplicates. The experiment was repeated on three consecutive days (Supplementary. Fig. S2). The kinetics of phage synthesis were similar in s-μg and 1-g conditions (Fig. 2A), and comparable to reactions in standard gravity and reaction tubes^9^. After two hours, plaques were detected in both conditions, and phage yield increased until 6 hours of incubation, after which phage yields plateaued. We observed a consistently higher number of phages in s-μg for all productive timepoints with the highest difference in PFU after three hours of incubation. At 3h, on average, three times as many PFU were counted in s-μg. Statistical analysis performed on all timepoints combined (unpaired t-test, p-value = 0.033) also confirmed a significantly higher number of synthesized phages in s-μg compared to 1-g (Supplementary. Fig. S3). We used transmission electron microscopy to confirm bacteriophage production. Typical icosahedral capsids of bacteriophage T7 were successfully visualized (Fig 1D). Notably, TEM samples with bacteriophages produced in s-μg contained a higher number of fully assembled phage capsids compared to 1-g samples. Conversely, fewer small, incomplete assemblies were visible in s-μg compared to 1-g (Supplementary. Fig. S4, S5).

**Figure 2.**
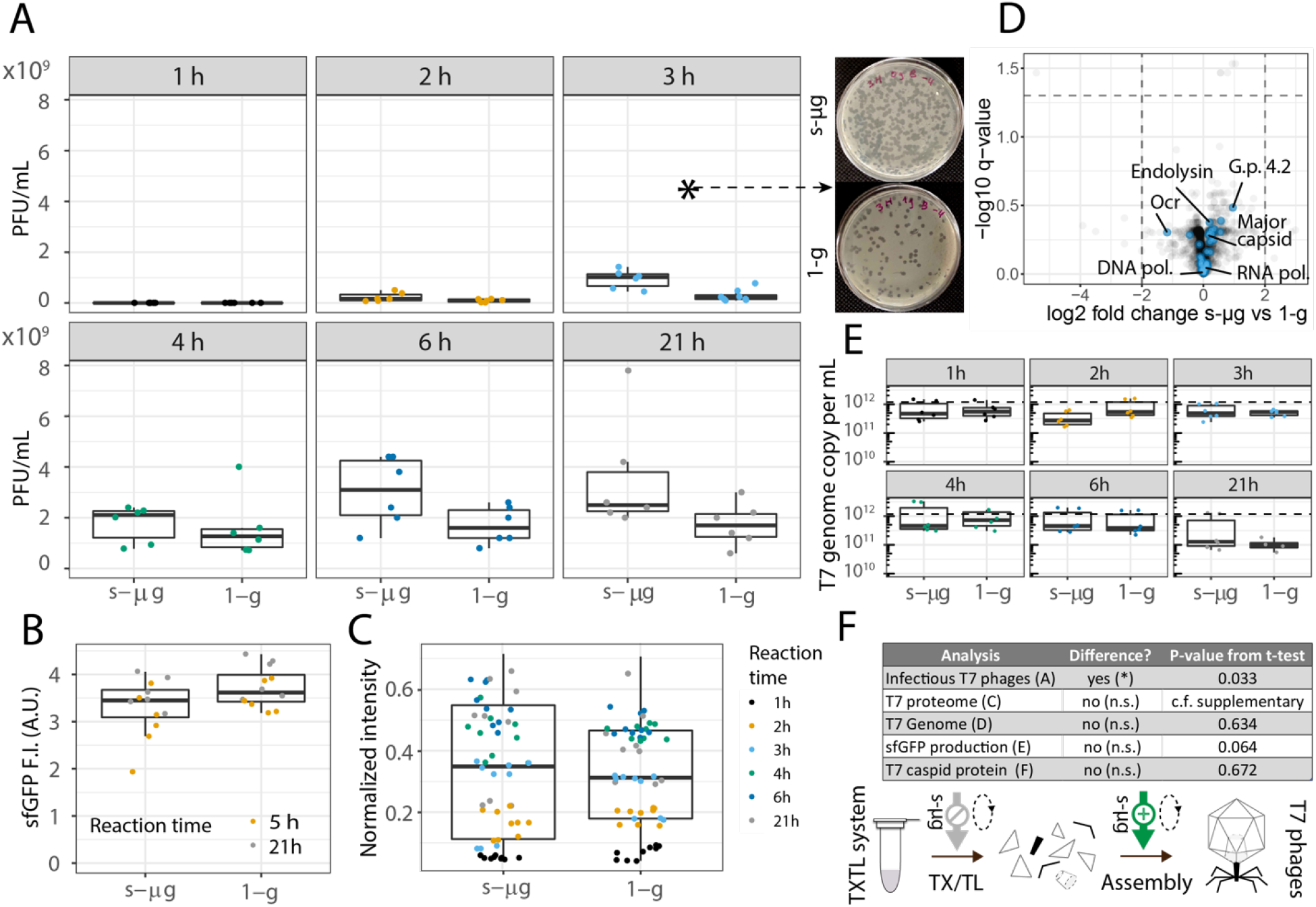
Quantitative analyses of T7 bacteriophage production in s-μg and 1-g conditions. A) Plaque forming units were determined at six different timepoints of the cell-free reaction. Images show example plaque assay plates from the 3 h timepoint. B) Production of sfGFP in control reactions set up in parallel to phage production. C) Relative quantification of the T7 major capsid protein by immuno-dot-blot. D) Proteomic analysis showed no significant differences in phage protein contents between the two conditions at 21 h. Volcano plot comparing s-μg against the 1-g condition. Gray dots represent the *E. coli* proteome and the blue dots correspond to T7 phage proteins with names of selected T7 proteins. E) Quantification of the T7 genome copy number by qRT-PCR shows no effect of s-μg on T7 genome copy number. The dashed line indicates the initial number of T7 DNA molecules in the samples (1.204×10^12^ per mL, 2 nM). F) Summary of the performed analyses, their results and statistical significance.

Since we measured an increased number of infectious phages in the s-μg environment, we next asked if the s-μg conditions enhanced synthesis reactions or phage assembly. A parallel TXTL reaction producing superfolder green fluorescent protein (sfGFP) did not show a significant difference in fluorescence intensity after 5 or 21 hours of incubation between s-μg and 1-g (Fig. 2B, unpaired t-test, p = 0.064). This suggests that the enhanced number of phages obtained in s-μg may not be the result of enhanced transcription or translation, but may rather be due to improved phage assembly. To support this hypothesis, we compared the composition of TXTL reactions from both conditions with several additional quantitative methods.

To gain insights in the phage protein production along the kinetics, we performed immuno-dot-blot assays targeting the major capsid protein, the most abundant structural T7 phage protein (Fig. 2C). Over the time course of the experiments, we did not find an overall significant difference (unpaired t-test, p-value = 0.672) in capsid protein signal between s-μg and 1-g conditions. To extend our analysis to the entire T7 proteome, the protein content of the 21 hours endpoint samples were analyzed by mass spectrometry. Over 1000 *E. coli* proteins were detected in each preparation and, as expected, showed similar composition in all samples (Supplementary. Fig. S6A), because our TXTL reagents consist of an *E. coli* lysate. In total, 80% of the bacteriophage T7 proteome was detected in s-μg and 1-g, including hypothetical (e.g. gene product 4.2), structural (e.g. major capsid protein) and non-structural proteins (e.g. DNA polymerase) (Supplementary. Fig. S6B). Using t-tests and q-values adjusted for false discovery rate, we did not observe significant differences between s-μg and 1-g for the 45 detected phage proteins (Fig. 2D, Supplementary. Table. 1). Hierarchical clustering of the detected T7 proteins did not lead to any visible patterns (Supplementary. Fig. S7). The proteomics results therefore support our previous findings that s-μg did not alter protein synthesis (Fig. 2F).

The bacteriophage T7 genome has been previously shown to replicate in TXTL reactions^26^. We hypothesized that the higher number of infectious phages in s-μg could stem from differences in genome replication or DNA degradation. To measure if s-μg samples contained higher amounts of DNA, we used quantitative real-time PCR (qRT-PCR) to track the bacteriophage T7 genome copy number over time^27^ (Fig. 2E). Similar to our protein content comparisons, no significant difference was observed in the DNA copy numbers between s-μg and 1-g conditions (one-way ANOVA, p-value = 0.223), indicating that the higher phage yield in s-μg was not a result of differences in DNA content. We observed no significant changes in the DNA concentration over time (one-way ANOVA, p-value = 0.0966), indicating no replication in s-μg or 1-g, possibly due to our high DNA starting concentration. Alternatively, replication and degradation of the linear DNA template may have been balanced.

In summary, our results show enhanced bacteriophage yields in simulated microgravity by 2D-clinorotation. Our quantitative analyses of protein and DNA contents revealed no differences between s-μg and 1-g samples. Since proteins and DNA are the building materials for virion assembly, we conclude that the increase in infectious bacteriophage particles is most likely due to enhanced self-assembly of bacteriophage T7 in simulated microgravity, as previously shown for protein crystal assembly in microgravity^14^. We hypothesize that the reduction of sedimentation in our experiments explains the improved self-assembly. Orbital shaking experiments also yielded increased PFU counts and support our hypothesis that preventing sedimentation and increased mixing improve bacteriophage production (Supplementary. Fig. S8). To enhance Earth-based cell-free production of other potential therapeutics that rely on self-assembly, it may be beneficial to prevent sedimentation by clinorotation or shaking. In conclusion, our results indicate that microgravity conditions might improve cell-free biomanufacturing yields, which is encouraging for the vision of an “astropharmacy” enabled by cell-free synthesis. As the next step, it should be tested how genuine spaceflight conditions, including increased radiation, influence cell-free production of self-assembled therapeutic agents.

## Materials and Methods

### TXTL extract preparation

TXTL systems preparation was adapted from Silverman et al.^28^, including adaptions described by Falgenhauer et al.^29^ *E. coli* BL21 Rosetta 2 were streaked overnight on an agar plate containing chloramphenicol. One colony was picked and inoculated overnight in 50 mL 2xYT supplemented with chloramphenicol for growth at 37°C. After a minimum of 15 hours, 20 mL of the stationary culture was used to inoculate 400 mL of 2xYT + P media (16 g/L tryptone, 10 g/L yeast extract, 5 g/L sodium chloride, 7 g/L potassium phosphate dibasic, 3 g/L potassium phosphate monobasic) in a 1 L baffled flask. Cells were grown at 40 °C and 200 RPM to 3.0 ± 0.2 OD_600_. Centrifuge bottles were filled up to 300 mL and centrifuged for 10 minutes at 4,000xg at 4 °C and supernatants were discarded. The pellets were washed three times with 25 mL buffer S30A (50 mM Tris-base, 14 mM Mg-glutamate, 60 mM K-glutamate, 2 mM DTT, brought to pH 7.7 with acetic acid). The washing steps were followed by a centrifugation step at 4,000xg at 4 °C for 10 minutes. A fourth centrifugation step at 3,000xg at 4 °C for 10 minutes enabled the removal of the remaining traces of buffer. The pellets were then resuspended in 1 mL of Buffer S30A per gram of pellet and supplemented with 0.5 mg/mL of lysozyme (from chicken egg, >40,000 units/mg, Sigma). The resuspended pellets were incubated for 10 minutes on ice. 1 mL of the suspension was aliquoted into 1.5 mL Eppendorf tubes. The pellet suspensions were then lysed with a sonicator (QSonica Q125 with a 3.175 mm diameter probe, 50% amplitude, 20 kHz, and 10 seconds ON/OFF pulses). Each sample was sonicated until reaching 250 J input. Using a 100 mM stock solution, 1 mM of DTT was added to each crude lysate immediately after sonication. The cell lysate was centrifuged for 10 minutes at 4 °C and 12,000xg. The supernatant was removed and placed into an incubator set up at 37 °C and 200 RPM for 80 minutes. After the run-off reaction, the supernatant was centrifuged for 10 minutes at 4 °C and 12,000xg. Finally, the extract was dialyzed for 3 hours against buffer S30B (50 mM Tris-base, 14 mM Mg-glutamate, 60 mM K-glutamate, 2 mM DTT, pH 8.2) in a 10k MWCO cassette (Thermofisher). Finally, the dialyzed extract was centrifuged for 10 minutes at 4 °C and 12,000xg. The supernatant was aliquoted, snap-frozen into liquid nitrogen, and stored at -80 °C.

### TXTL mastermix reaction

The final TXTL reaction mixture is composed of the following reagents: 33% v/v of *E. coli* extract, 10 mM ammonium glutamate; 1.2 mM ATP; 0.850 mM each of GTP, UTP, and CTP; 0.034 mg/mL folinic acid; 0.175 mg/mL yeast tRNA; 2 mM amino acids; 30 mM 3-PGA; 0.33 mM NAD; 0.27 mM CoA; 1 mM putrescine; 1.5 mM spermidine; 57 mM HEPES, 3.5% PEG 8000, 5 μM of Chi6 linear DNA. Mg-glutamate and K-glutamate were optimized for phage production and set respectively to 7 mM and 170 mM. T7 Phage genomic DNA (Bioron) was set to 2 nM (1.204×10^12^ DNA molecules per mL).

### Clinostat operation and TXTL loading

The 2D-clinostat used in this study was developed and kindly provided by the DLR Institute of Aerospace Medicine, Department of Gravitational Biology and has been used in previous studies on the effects of microgravity on biological systems^22,24^. The clinorotation speed was 60 RPM. The TXTL mastermix was prepared with all components except T7 Phage genomic DNA. The mastermix was split into 20-μL aliquots, and each was introduced inside its own PTFE tube (1/16”ID x 1/8”OD). The PTFE tubes were sealed with metal caps (Fig. 1C). The T7 Phage genomic DNA was added just before loading the PTFE tubes, to minimize the reaction time before the experiment. Instead of T7 DNA, negative controls contained nuclease-free water, and translation control tubes contained a plasmid expressing sfGFP under the control of a strong constitutive promoter. For the samples analyzed with proteomics, the loaded volume was 50 μL. The PTFE tubes were inserted into hollow PVC tubes fitting the clinostat rotational tray. The clinostat containing all the tubes was held in an incubator at 29 °C. The samples of TXTL were taken after 1 h, 2 h, 3 h, 4 h, 6 h, and 21 h of incubation. For each timepoint, plaque assays were performed in duplicate with serial dilutions by plating samples immediately.

### Phage infectivity assay

At each timepoint, serial decimal dilutions of the TXTL reactions were prepared in Luria-Bertani (LB) medium. Then, 5 μL of each serial dilution was mixed with 130 μL of *E. coli* DSM 613 (OD_600_ between 0.8 and 1) and incubated for 3 min at 37 °C. All 135 μL were transferred into 1.75 mL of soft LB medium, mixed well, and poured onto a 60 mm petri dish. Each petri dish was left for 30 min at room temperature to solidify and then transferred to 37 °C. The plaques were counted after 3 h of incubation. After preparing the serial dilutions for the infectivity assays, samples were frozen at -80 °C with 25% (v/v) glycerol until used for other analysis of protein content, replication, and assembly.

### sfGFP fluorescence

sfGFP fluorescence was measured after 5 and 21 h of incubation. The TXTL reactions contained a plasmid expressing sfGFP under the control of a strong constitutive *E. coli* promoter. The fluorescence intensity (excitation: 485 nm; emission: 515 nm) was measured with a plate-reader (TECAN Infinite M200 Pro) and compared between s-μg and 1-g samples (10 μL of the reaction sample were used).

### Transmission Electron Microscopy

30 μL of the diluted TXTL samples from the 21 h timepoint was further diluted to 200 μL with TBS (Tris 20mM, NaCl 150 mM, pH 7.6). The samples were extracted by adding 10 μL of chloroform and centrifuged for 10 minutes at 13,000xg. The upper aqueous phase was transferred to a new tube. The bacteriophages were precipitated by adding 50 μL of PEG/NaCl 5x buffer (PEG-8000 20%, NaCl 2.5 M). The tubes were incubated at 4 °C overnight. The bacteriophages were pelleted by centrifugation for 10 minutes at 13,000xg. The bulks of supernatants were removed. The samples were centrifuged again for 10 minutes at 13,000xg and the supernatant was completely removed. The pellets were resuspended in 20 μL TBS. Carbon-coated copper grids (400 mesh) were hydrophilized by glow discharging (PELCO easiGlow, Ted Pella, USA). 5 μL of the primary antibody solution (T7 tag, PA1-32386, Thermofisher) was applied onto the hydrophilized grids. After 2 minutes of incubation, the solutions were briefly blotted and 5 μL of the samples were applied onto the grids. Alternatively, for Fig. S4 and Fig. S5, “antibody coated” grids were incubated in diluted sample solution (1:40) for 1 hour under shaking. After two short washing steps in droplets of double distilled water, samples were stained with 2% uranyl acetate. Samples were analyzed with a JEOL JEM-2100 transmission electron microscope using an acceleration voltage of 120 kV. Images were acquired with a F214 FastScan CCD camera (TVIPS, Gauting).

### Proteomics

Proteins contained in TXTL samples (50 μL) were reduced by adding 5 mM Tris(2-caboxyethyl)phosphine at 90 °C for 15 minutes, followed by alkylation (10mM iodoacetamide, 30 min at 25 °C). The amount of extracted proteins was measured using BCA protein assay (Thermo Fisher Scientific). 50 μg total protein was then digested with 1 μg trypsin (Promega) overnight at 30 °C in the presence of 0.5% SLS. Following digestion, SLS was precipitated with trifluoroacetic acid (TFA, 1.5% final concentration) and peptides were purified using Chromabond C18 microspin columns (Macherey-Nagel). Acidified peptides were loaded on spin columns equilibrated with 400 μL acetonitrile and then 400 μL 0.15% TFA. After peptide loading, a washing step with 0.15% TFA was performed, followed by elution using 400 μL 50% acetonitrile. Eluted peptides were then dried by vacuum concentrator and reconstituted in 0.15% TFA.

Peptide mixtures were analyzed using liquid chromatography-mass spectrometry carried out on an Exploris 480 instrument connected to an Ultimate 3000 RSLC nano with a Prowflow upgrade and a nanospray flex ion source (all Thermo Scientific). Peptide separation was performed on a reverse phase HPLC column (75 μm x 42 cm) packed in-house with C18 resin (2.4 μm, Dr. Maisch). The following separating gradient was used: 94% solvent A (0.15% formic acid) and 6% solvent B (99.85% acetonitrile, 0.15% formic acid) to 25% solvent B over 40 minutes and to 35% B for additional 20 minutes at a flow rate of 300 mL/min. DIA-MS acquisition method was adapted from D. Bekker-Jensen et al.^30^ In short, Spray voltage were set to 2.0 kV, funnel RF level at 55, and heated capillary temperature at 275 °C. For DIA experiments full MS resolutions were set to 120.000 at m/z 200 and full MS AGC target was 300% with an IT of 50 ms. Mass range was set to 350–1,400. AGC target value for fragment spectra was set at 3,000%. 45 windows of 15 Da were used with an overlap of 1 Da. Resolution was set to 15,000 and IT to 22 ms. Stepped HCD collision energy of 25, 27.5, 30% was used. MS1 data was acquired in profile, MS2 DIA data in centroid mode. Analysis of DIA data was performed using DIA-NN version 1.8^31^ using Uniprot protein databases from *E. coli* and bacteriophage T7. Full tryptic digest was allowed with two missed cleavage sites, and oxidized methionines and carbamidomethylated cysteines. Match between runs and remove likely interferences were enabled. The neural network classifier was set to the single-pass mode, and protein inference was based on genes. Quantification strategy was set to any LC (high accuracy). Cross-run normalization was set to RT-dependent. Library generation was set to smart profiling. DIA-NN exports were further statistically evaluated using a modified SafeQuant script made compatible to process DIA-NN data.

### Dot-Blot

2 μL of the diluted TXTL samples were blotted on a nitrocellulose membrane and let dry for 15 minutes. The membrane was incubated for 1 hour with blocking buffer (TBST 1x, 5% w/v non-fat dry milk) on a shaker at RT. The membrane was rinsed 2 times with water and incubated for 1 hour on a shaker at room temperature (RT) with the primary antibody (T7 tag, PA1-32386, Thermofisher) diluted in the blocking buffer (1:10,000). The membrane was washed 3 times with the wash buffer (TBST 1x) and incubated for 1 hour on a shaker at RT with the secondary antibody (IRDye® 800CW Goat anti-Rabbit IgG, LI-COR) diluted (1:20,000) in the blocking buffer. Finally, the membrane was washed 3 times with the wash buffer and scanned for fluorescence with an LI-COR Odyssey DLx. The calibration curve showed that the concentration of synthesized phages from the experiment was in the linear range of detection (Supplementary. Fig. S8). Each experimental day was blotted on separated membranes and normalized to a standard.

### qRT-PCR

To determine the bacteriophage T7 DNA copy number, qRT-PCR was performed. Each TXTL sample was diluted 100x in nuclease-free water, and 5 μL of the diluted TXTL was used for qRT-PCR, as described earlier for T7 phage produced in TXTL^32^. The assays were performed in technical duplicates for each of the timepoints from all three experimental replicates. The primers and the program were adapted from Peng et al.^27^ (Supplementary. Table S2, Supplementary. Table S3). The kit used for the qRT-PCR reaction was NEB Luna® Universal qPCR & qRT-PCR. As a standard, 5 μL of T7 Phage DNA (Bioron) was used. The standard was diluted in nuclease-free water. Different dilutions were used: 0.2 nM (610,000,000 T7 DNA molecules), 0.2×10^−2^ nM (6,100,000 T7 DNA molecules), 0.2×10^−4^ nM (61,000 T7 DNA molecules), and 0.2×10^−6^ nM (610 T7 DNA molecules). The determined DNA copy number in each sample was corrected for the original TXTL reaction concentration and volume (100x concentrated, 20 μL).

## Acknowledgements

This work was supported by Deutsche Forschungsgemeinschaft grant NI 2040/1-1. Ralf Moeller and Bruno Pavletic received funding from the following DLR (German Aerospace Center) grants: ISS LIFE (Program RF-FuW, Teilprogramm: 475) and GANDALF PhD Graduate School and by an ERASMUS+ fellowship (2019-1-HR01-KA103-060250 for Bruno Pavletic). These results will be included in the PhD thesis of Bruno Pavletic. We thank Horst Henseling for constructing the clinostat sample holders, Nadiia Pozhydaieva and Katharina Höfer for advice on phage handling, Jing Yuan and Shan Jiang for advice on immuno-dot-blot, Laura Czech for advice on TEM sample preparation, Ruth Hemmersbach, Jens Hauslage, and Kai Wasser for the 2-D clinostat and for advice on clinorotation, and Anke Becker for discussion of the project idea. The authors express their gratitude to the entire ESA Topical Team Space Synthetic Biology (Contract No. 4000132667/20/NL/CLR/pt).

## Data availability

The authors declare that all relevant data supporting the findings of this study are available within the paper and its Supplementary information files. Additional data are available from the corresponding author upon request.

## References

1. Patel, Z. S. et al. Red risks for a journey to the red planet: The highest priority human health risks for a mission to Mars. Npj Microgravity 6, 1–13 (2020).

2. Crucian, B. E. et al. Immune System Dysregulation During Spaceflight: Potential Countermeasures for Deep Space Exploration Missions. Front. Immunol. 9, (2018).

3. Snyder, J. E., Walsh, D., Carr, P. A. & Rothschild, L. J. A Makerspace for Life Support Systems in Space. Trends Biotechnol. 37, 1164–1174 (2019).

4. Stark, J. C. et al. On-demand biomanufacturing of protective conjugate vaccines. Sci. Adv. 7, eabe9444 (2021).

5. Wilding, K. M. et al. Endotoxin-Free E. coli-Based Cell-Free Protein Synthesis: Pre-Expression Endotoxin Removal Approaches for on-Demand Cancer Therapeutic Production. Biotechnol. J. 14, 1800271 (2019).

6. Nagappa, L. K. et al. A ubiquitous amino acid source for prokaryotic and eukaryotic cell-free transcription-translation systems. Front. Bioeng. Biotechnol. 10, 992708–992708 (2022).

7. Rummel, J. D. From Planetary Quarantine to Planetary Protection: A NASA and International Story. Astrobiology 19, 624–627 (2019).

8. Emslander, Q. et al. Cell-free production of personalized therapeutic phages targeting multidrug-resistant bacteria. Cell Chem. Biol. 29, 1434-1445.e7 (2022).

9. Rustad, M., Eastlund, A., Jardine, P. & Noireaux, V. Cell-free TXTL synthesis of infectious bacteriophage T4 in a single test tube reaction. Synth. Biol. 3, (2018).

10. Bundy, B. C., Franciszkowicz, M. J. & Swartz, J. R. Escherichia coli-based cell-free synthesis of virus-like particles. Biotechnol. Bioeng. 100, 28–37 (2008).

11. Brives, C. & Pourraz, J. Phage therapy as a potential solution in the fight against AMR: obstacles and possible futures. Palgrave Commun. 6, 1–11 (2020).

12. Ando, H., Lemire, S., Pires, D. P. & Lu, T. K. Engineering Modular Viral Scaffolds for Targeted Bacterial Population Editing. Cell Syst. 1, 187–196 (2015).

13. Nooraei, S. et al. Virus-like particles: preparation, immunogenicity and their roles as nanovaccines and drug nanocarriers. J. Nanobiotechnology 19, 59 (2021).

14. McPherson, A. & DeLucas, L. J. Microgravity protein crystallization. NPJ Microgravity 1, 15010 (2015).

15. Bell, D. et al. Self-Assembly of Protein Fibrils in Microgravity. Gravitational Space Res. 6, 10–26 (2018).

16. Zhou, J. et al. Effects of sedimentation, microgravity, hydrodynamic mixing and air–water interface on α-synuclein amyloid formation. Chem Sci 11, 3687–3693 (2020).

17. Chang, D., Paulsen, A., Johnson, T. C. & Consigli, R. A. Virus protein assembly in microgravity. Adv. Space Res. 13, 251–257 (1993).

18. Koszelak, S., Day, J., Leja, C., Cudney, R. & McPherson, A. Protein and virus crystal growth on international microgravity laboratory-2. Biophys. J. 69, 13–19 (1995).

19. Acres, J. M., Youngapelian, M. J. & Nadeau, J. The influence of spaceflight and simulated microgravity on bacterial motility and chemotaxis. Npj Microgravity 7, 1–11 (2021).

20. Rosenstein, A. H. & Walker, V. K. Fidelity of a Bacterial DNA Polymerase in Microgravity, a Model for Human Health in Space. Front. Cell Dev. Biol. 9, (2021).

21. Eiermann, P. et al. Adaptation of a 2-D Clinostat for Simulated Microgravity Experiments with Adherent Cells. Microgravity Sci. Technol. 25, 153–159 (2013).

22. Brungs, S., Kolanus, W. & Hemmersbach, R. Syk phosphorylation – a gravisensitive step in macrophage signalling. Cell Commun. Signal. 13, 9 (2015).

23. Klaus, D. M., Todd, P. & Schatz, A. Functional weightlessness during clinorotation of cell suspensions. Adv. Space Res. Off. J. Comm. Space Res. COSPAR 21, 1315–1318 (1998).

24. Trotter, B. et al. The influence of simulated microgravity on the proteome of Daphnia magna. Npj Microgravity 1, 1–10 (2015).

25. Kemp, P., Garcia, L. R. & Molineux, I. J. Changes in bacteriophage T7 virion structure at the initiation of infection. Virology 340, 307–317 (2005).

26. Shin, J., Jardine, P. & Noireaux, V. Genome Replication, Synthesis, and Assembly of the Bacteriophage T7 in a Single Cell-Free Reaction. ACS Synth. Biol. 1, 408–413 (2012).

27. Peng, X., Leal, J., Mohanty, R., Soto, M. & Ghosh, D. Quantitative PCR of T7 Bacteriophage from Biopanning. JoVE J. Vis. Exp. e58165 (2018) doi:10.3791/58165.

28. Silverman, A. D., Kelley-Loughnane, N., Lucks, J. B. & Jewett, M. C. Deconstructing Cell-Free Extract Preparation for in Vitro Activation of Transcriptional Genetic Circuitry. ACS Synth. Biol. 8, 403–414 (2019).

29. Falgenhauer, E. et al. Evaluation of an E. coli Cell Extract Prepared by Lysozyme-Assisted Sonication via Gene Expression, Phage Assembly and Proteomics. ChemBioChem 22, 2805–2813 (2021).

30. Bekker-Jensen, D. B. et al. A Compact Quadrupole-Orbitrap Mass Spectrometer with FAIMS Interface Improves Proteome Coverage in Short LC Gradients. Mol. Cell. Proteomics MCP 19, 716–729 (2020).

31. Demichev, V., Messner, C. B., Vernardis, S. I., Lilley, K. S. & Ralser, M. DIA-NN: neural networks and interference correction enable deep proteome coverage in high throughput. Nat. Methods 17, 41–44 (2020).

32. Vogele, K., Falgenhauer, E., von Schönberg, S., Simmel, F. C. & Pirzer, T. Small Antisense DNA-Based Gene Silencing Enables Cell-Free Bacteriophage Manipulation and Genome Replication. ACS Synth. Biol. 10, 459–465 (2021).

