## Supplementary information for "Enhanced assembly of bacteriophage T7 produced in cell-free reactions under simulated microgravity"

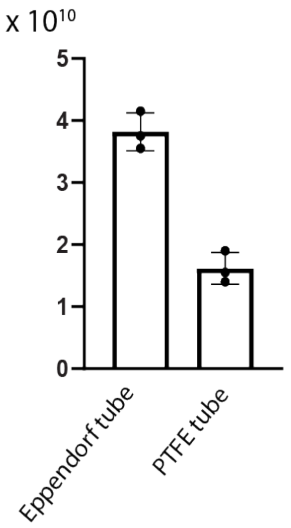


**Supplementary Figure S1**. Cell-free synthesis efficiency of T7 bacteriophages decreases in PTFE tubes. Plaque assay variability remains low when PTFE tubes are used as a reaction vessel.

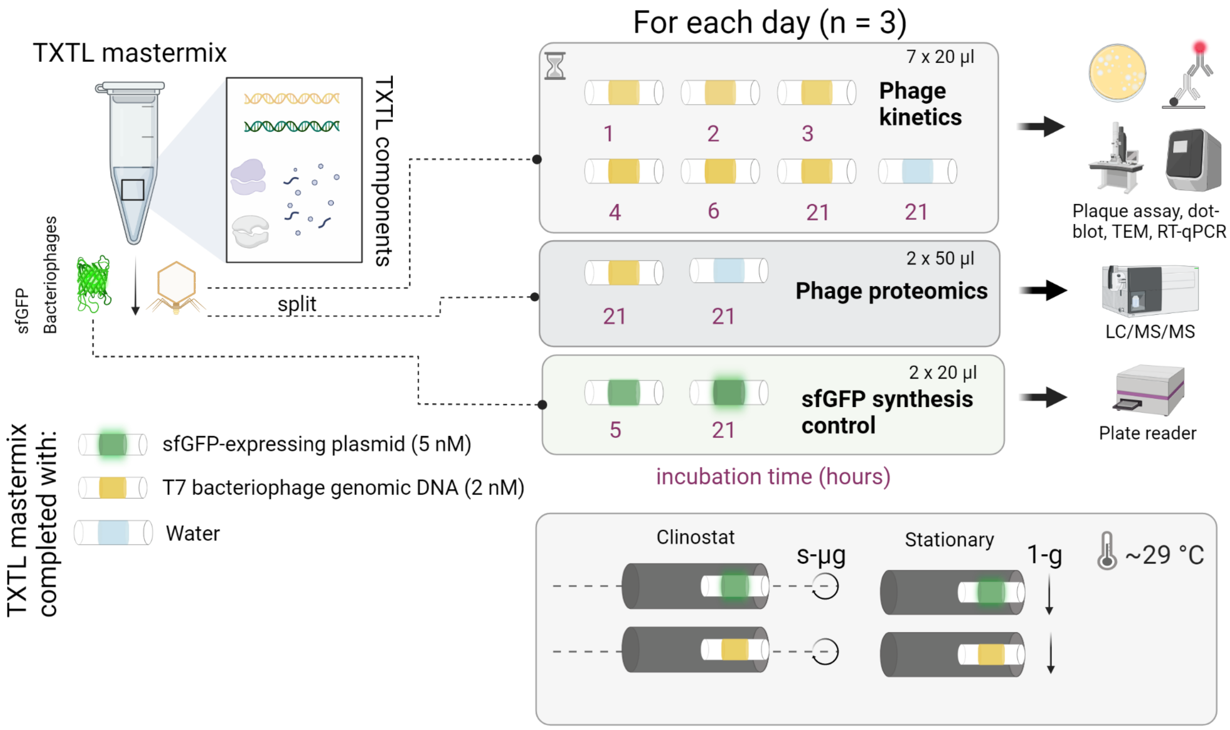


**Supplementary Figure S2**. Experimental set-up of the cell-free synthesis operated in simulated microgravity and in stationary control conditions. The experimental set-up shown corresponds to one day of experiment and was repeated 3 times.

A

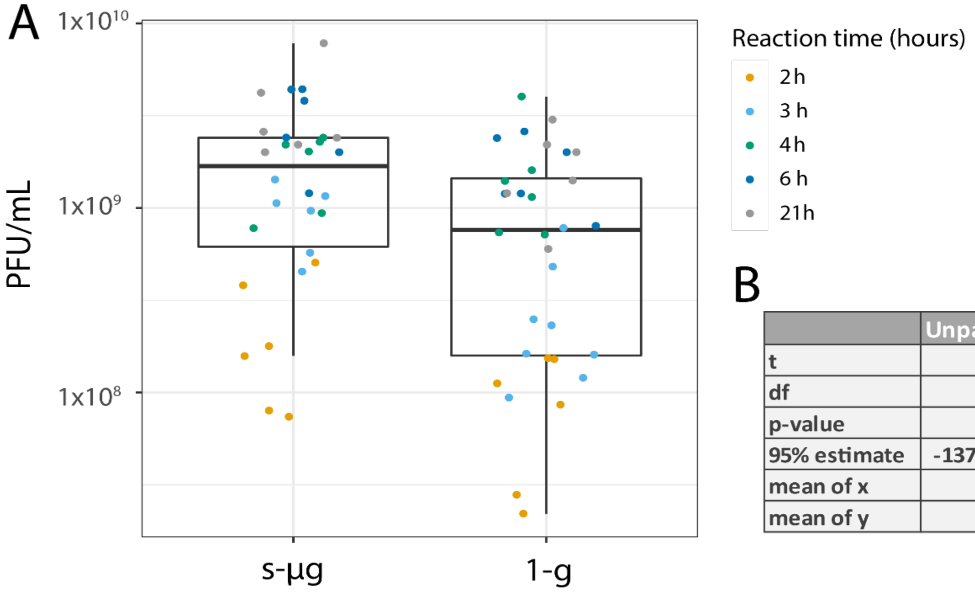


B

| | Unpaired t-test |
|---|---|
| t | -2.1862 |
| df | 55.887 |
| p-value | 0.03301 |
| 95% estimate | -1370003507 / -59797673 |
| mean of x | 869210526 |
| mean of y | 1584111117 |

**Supplementary Figure S3.** A) Summary PFU/mL of bacteriophages T7 synthesized in s-µg and 1-g. B) Statistical summary table for unpaired t-test comparing the two conditions.

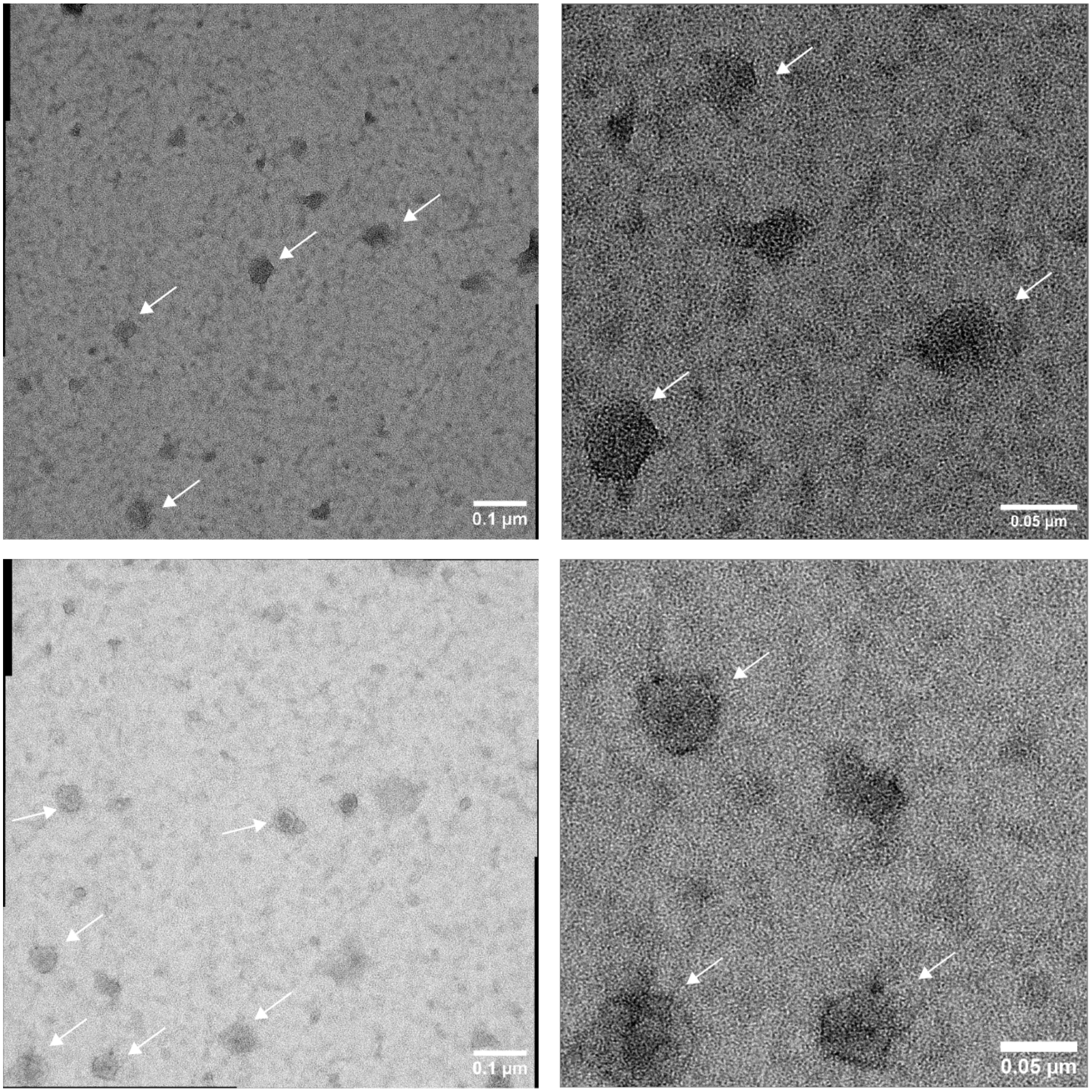


**Supplementary Figure S4.** T7 bacteriophages synthesized in s-µg visualized by transmission electron microscopy. White arrows indicate presumed fully assembled phage capsids.

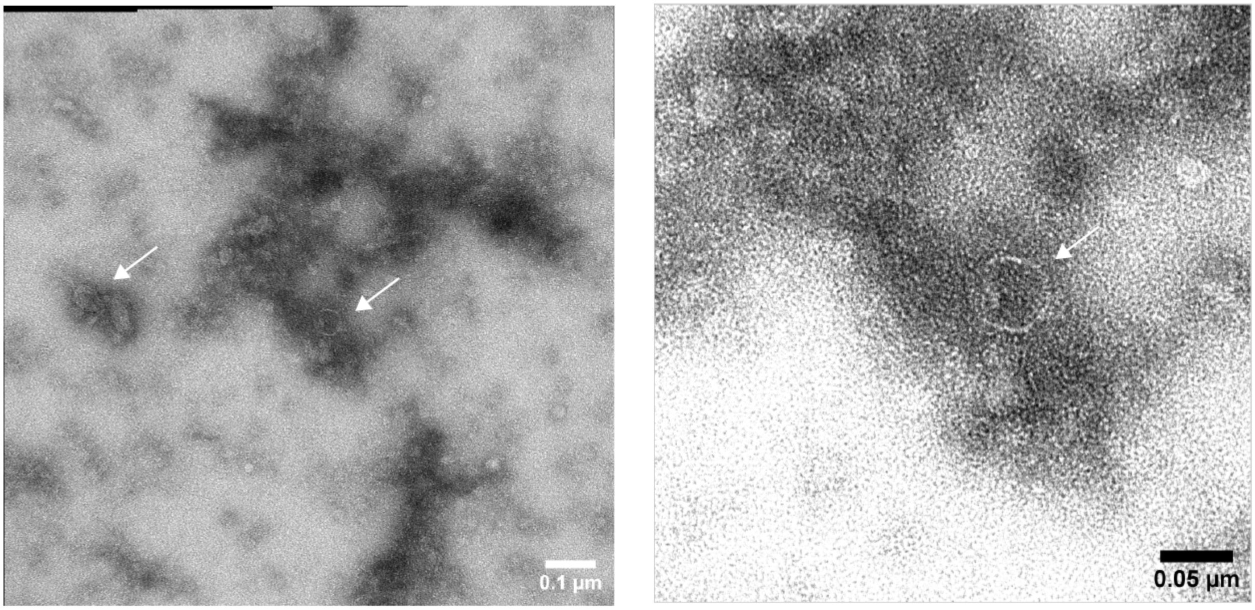


**Supplementary Figure S5**. T7 bacteriophages synthesized in 1-g visualized by transmission electron microscopy. White arrows indicate presumed fully assembled phage capsids.

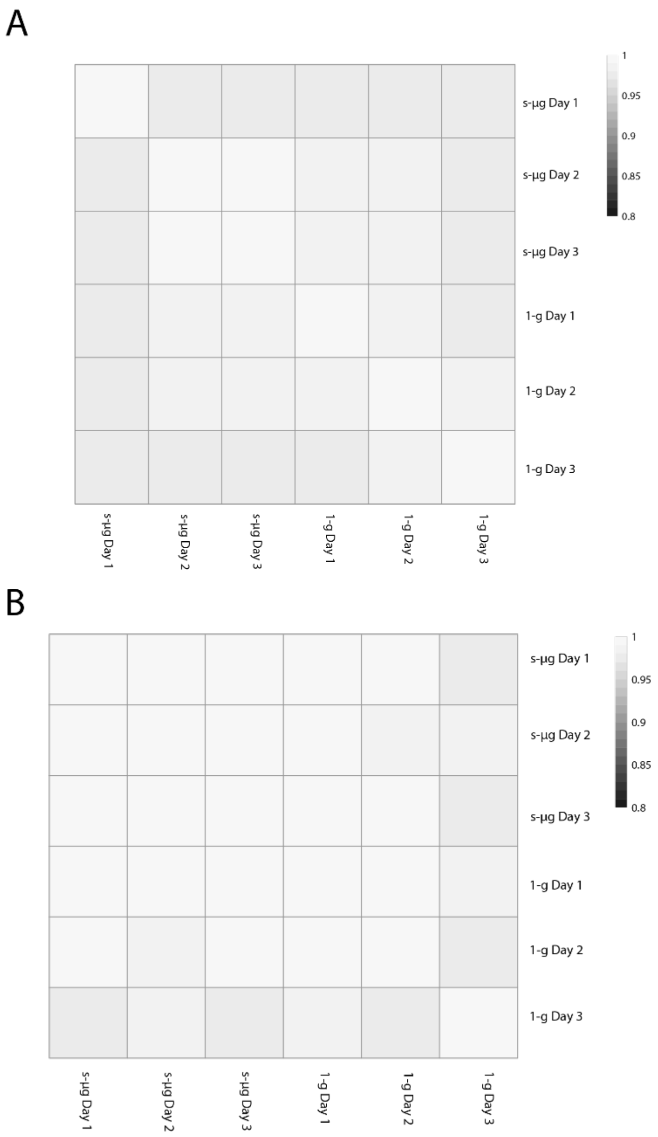


**Supplementary Figure S6**. Pearson correlation matrix of the detected *E. coli* proteome (A) and T7 proteins (B) between the three experimental replicates in s-μg and 1-g.

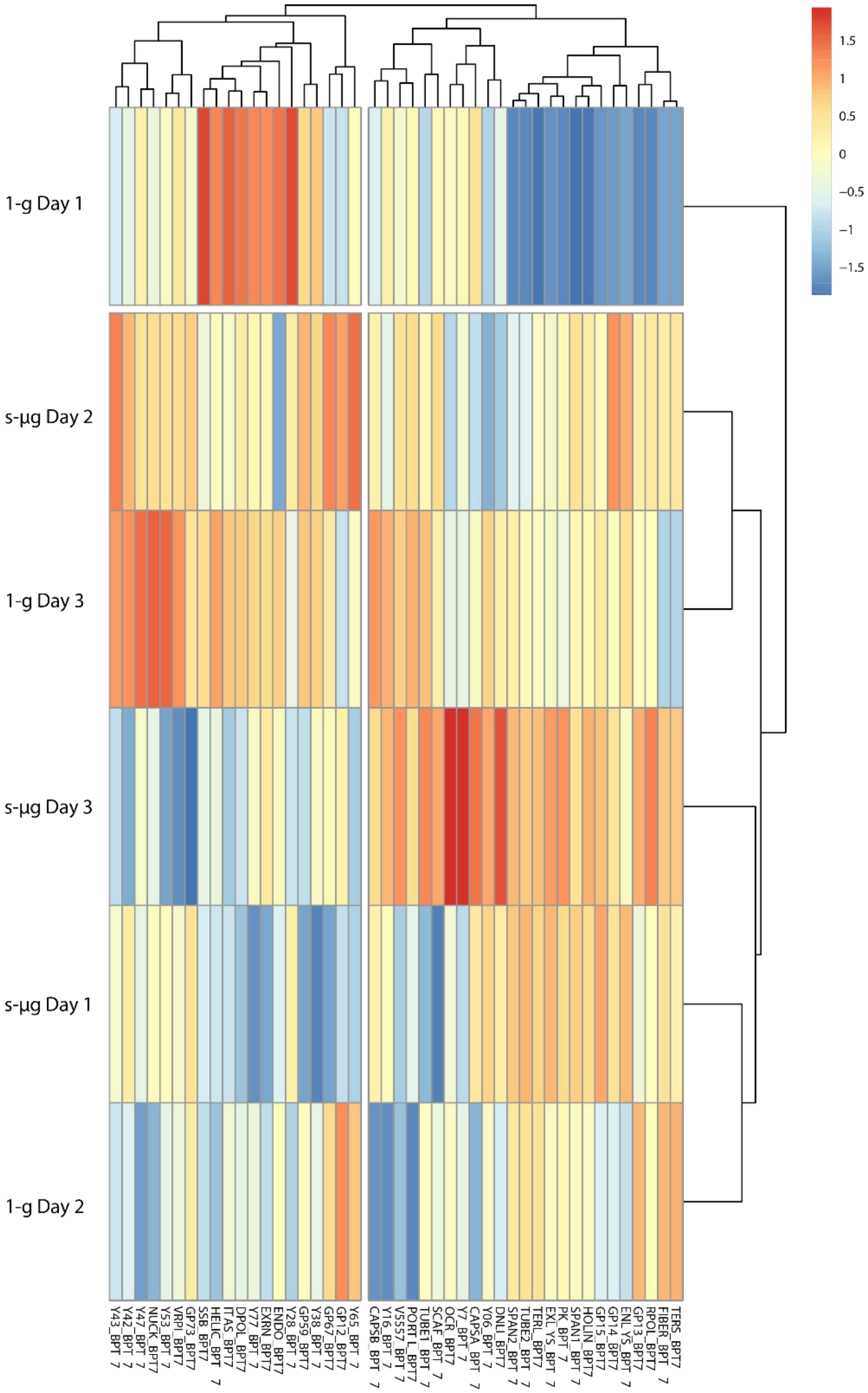


**Supplementary Figure S7.** Heatmap representation of the T7 bacteriophage proteins detected in 1-g and s-µg. Vertical columns represent the proteins identified by LC-MS/MS. Rows depict the biological replicates in s-µg and 1-g. The protein intensities values were centered and scaled in column direction and used for hierarchical clustering of the samples by using Euclidian distance and average method (constructed using the pheatmap package from R).

A

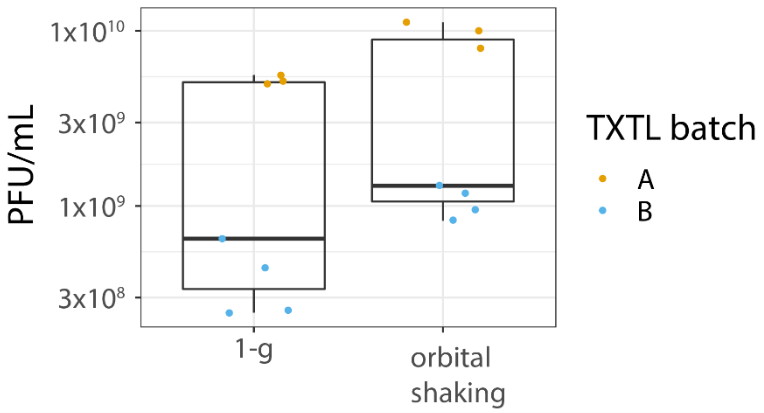


B

| | Unpaired t-test |
|---|---|
| t | -2.3384 |
| df | 13.124 |
| p-value | 0.03583 |
| 95% estimate | -9578256189 / -383629525 |
| mean of x | 2484857143 |
| mean of y | 7465800000 |

**Supplementary Figure S8**. A) Bacteriophage T7 cell-free synthesis comparison between 1-g conditions and orbital shaking at 60 RPM. Two different TXTL batches were used for this experiment. Batch A corresponds to the same batch used for the clinorotation experiment. Batch B is a different batch, which explains the differences in PFU yield. B) Statistical summary table for the shaking control experiment.

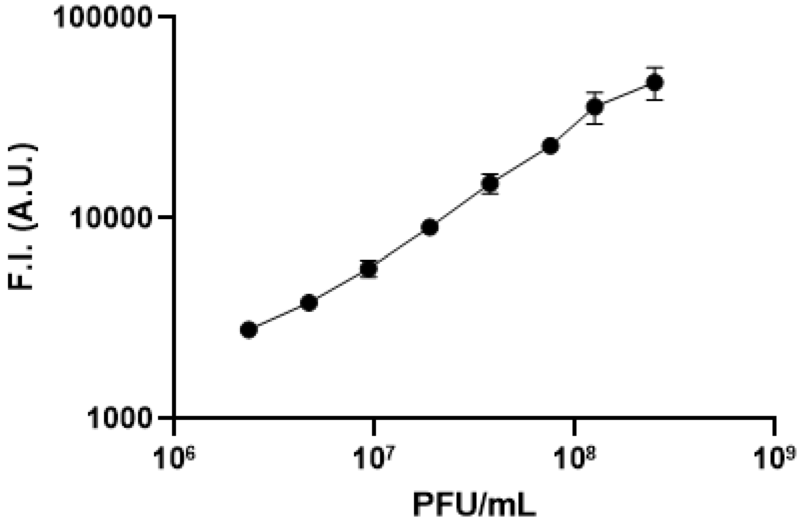


**Supplementary Figure S9.** Calibration curve for the dot-blot experiments. Pre-determined concentrations of T7 bacteriophages produced in TXTL systems were used to establish optimal scanning settings for the PFU/mL measured in the synthesis kinetics.

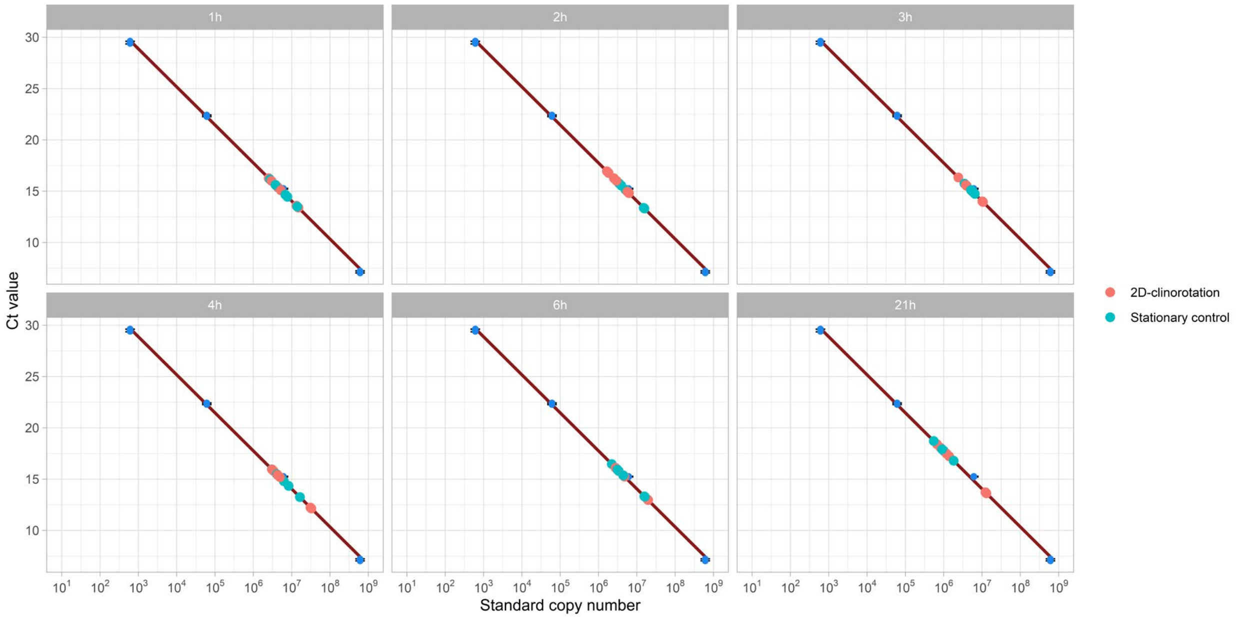


**Supplementary Figure S10. The RT-qPCR results per timepoint projected onto the standard curve.** Plotted are the total copy numbers in 5 µL of 100x diluted TXTL samples. The results were then corrected to 20 µL of the total reaction from which the DNA copy number per mL was determined.

**Supplementary Table S1.** List of T7 phage proteins detected by mass spectrometry and their associated statistics. Log2ratio, p-Value and q-Value correspond to the difference of averaged intensities peaks of the samples in s-µg compared to 1-g. The colormap of log2ratio is centered on 0 (negative values are red and positive values are blue).

| Protein name | Abbrevation | Protein description | log2ratio T7 s-µg | p-Value T7 s-µg | q-Value T7 s-µg |
|---|---|---|---|---|---|
| P03726 | EXLYS_BPT7 | Peptidoglycan transglycosylase gp16 | 0.09410517 | 0.619589159 | 0.829362696 |
| P03725 | GP15_BPT7 | Internal virion protein gp15 | 0.211401056 | 0.237367394 | 0.583862348 |
| P00573 | RPOL_BPT7 | T7 RNA polymerase | 0.059783722 | 0.75954832 | 0.898736323 |
| P00581 | DPOL_BPT7 | DNA-directed DNA polymerase | -0.00902532 | 0.922500358 | 0.970855339 |
| P00969 | DNLI_BPT7 | DNA ligase | -0.049660159 | 0.685584248 | 0.854089292 |
| P03748 | FIBER_BPT7 | Tail fiber protein | -0.042447949 | 0.863543454 | 0.944713949 |
| P03747 | TUBE2_BPT7 | Tail tubular protein gp12 | 0.051979995 | 0.730727255 | 0.882677549 |
| P03696 | SSB_BPT7 | Single-stranded DNA-binding protein | -0.033412415 | 0.765950785 | 0.902556252 |
| P03692 | HELIC_BPT7 | DNA helicase/primase | 0.034877351 | 0.820116661 | 0.933229546 |
| P03787 | V5557_BPT7 | Fusion protein 5.5/5.7 | 0.015180704 | 0.905155552 | 0.963883461 |
| P00806 | ENLYS_BPT7 | Endolysin | 0.204159787 | 0.019846392 | 0.420239352 |
| P03728 | PORTL_BPT7 | Portal protein | 0.108383002 | 0.403621646 | 0.686987295 |
| P03694 | TERL_BPT7 | Terminase, large subunit | 0.120603168 | 0.656456394 | 0.841631632 |
| P03786 | Y47_BPT7 | Protein 4.7 | 0.274465461 | 0.203623083 | 0.551173505 |
| P00513 | PK_BPT7 | Protein kinase 0.7 | 0.091480446 | 0.745839376 | 0.89052877 |
| P03716 | SCAF_BPT7 | Capsid assembly scaffolding protein | -0.107561677 | 0.497981473 | 0.747876316 |
| P00638 | EXRN_BPT7 | Exonuclease | -0.050901179 | 0.582761387 | 0.802891979 |
| P03724 | GP14_BPT7 | Internal virion protein gp14 | 0.189530578 | 0.137178708 | 0.50423504 |
| P03781 | NUCK_BPT7 | Nucleotide kinase gp1.7 | 0.351786207 | 0.038665058 | 0.47079694 |
| P03797 | Y38_BPT7 | Protein 3.8 | -0.033445295 | 0.796411635 | 0.921129341 |
| P03785 | ITAS_BPT7 | Inhibitor of toxin/antitoxin system | -0.024237527 | 0.899720733 | 0.960540744 |
| P03693 | TERS_BPT7 | Terminase, small subunit gp18 | -0.06368776 | 0.828420896 | 0.936881863 |
| P03793 | Y16_BPT7 | Protein 1.6 | 0.092167856 | 0.670571337 | 0.849237179 |
| P03750 | Y7_BPT7 | Protein 7 | -0.444467689 | 0.169306805 | 0.517799877 |
| P00641 | ENDO_BPT7 | Endonuclease I | -0.136896019 | 0.283079282 | 0.608539939 |
| P03704 | VRPI_BPT7 | Bacterial RNA polymerase inhibitor | 0.295345968 | 0.149202798 | 0.510033585 |
| P03800 | Y65_BPT7 | Protein 6.5 | 0.053760279 | 0.829980386 | 0.936881863 |
| P03796 | Y77_BPT7 | Protein 7.7 | -0.072891012 | 0.526687254 | 0.768103799 |
| P03746 | TUBE1_BPT7 | Tail tubular protein gp11 | -0.013333543 | 0.835764217 | 0.936881863 |
| P19726 | CAPSA_BPT7 | Major capsid protein | -0.084937177 | 0.688752183 | 0.855459168 |
| P03723 | GP13_BPT7 | Probable scaffold protein gp13 | -0.013554018 | 0.913590507 | 0.967689824 |
| P03803 | SPAN1_BPT7 | Spanin, inner membrane subunit | 0.329015148 | 0.220874041 | 0.566006835 |
| P03795 | Y28_BPT7 | Protein 2.8 | 0.015069037 | 0.92989573 | 0.974006114 |
| P20406 | GP59_BPT7 | Probable RecBCD inhibitor gp5.9 | 0.015859888 | 0.844823203 | 0.939308852 |
| P03784 | Y43_BPT7 | Protein 4.3 | 0.568749349 | 0.01470882 | 0.409374313 |
| P03798 | Y53_BPT7 | Protein 5.3 | 0.246981903 | 0.061591431 | 0.4787412 |
| P03751 | GP73_BPT7 | Protein 7.3 | 0.553323332 | 0.105829791 | 0.493904652 |
| P03780 | GP12_BPT7 | Inhibitor of dGTPase | -0.090927084 | 0.620417839 | 0.829362696 |
| P03802 | HOLIN_BPT7 | Holin | 0.224738299 | 0.413478971 | 0.691585669 |
| P03788 | SPAN2_BPT7 | Spanin, outer lipoprotein subunit | 0.142527365 | 0.72600417 | 0.8805728 |
| P03778 | Y06_BPT7 | Protein 0.6B | 0.02362296 | 0.961529344 | 0.981947004 |
| P03783 | Y42_BPT7 | Preotein 4.2 | 0.957644976 | 0.006037242 | 0.327878033 |
| P19727 | CAPSB_BPT7 | Minor capsid protein | 0.164073009 | 0.174660113 | 0.518216805 |
| P03801 | GP67_BPT7 | Protein 6.7 | 0.015852896 | 0.946346575 | 0.975595572 |
| P03775 | OCR_BPT7 | Protein Ocr | -1.177699875 | 0.122367158 | 0.497137089 |

**Supplementary Table S2.** Contents of each RT-qPCR reaction.

| Component | Volume |
|---|---|
| Luna® Universal qPCR Master Mix (2x) | 10 µL |
| Forward primer (CCTCTTGGGAGGAAGAGATTTG, 10 nM) | 2.5 µL |
| Reverse primer (TACGGGTCTCGTAGGACTTAAT, 10 nM) | 2.5 µL |
| TXTL sample (100x diluted in nuclease-free water) | 5 µL |

**Supplementary Table S3.** The program used for the RT-qPCR to detect bacteriophage T7 genomic DNA in TXTL.

| Temperature | Time | No. Cycles |
|---|---|---|
| **50 °C** | 120 s | 1x |
| **95 °C** | 120 s | 1x |
| **95 °C** | 15 s | 40x |
| **60 °C** | 60 s | |
| **95 °C** | 15 s | 1x |
| **60 °C** | 60 s | 1x |
| **95 °C** | 15 s | 1x |
